# Non-muscle myosin 2 filaments are processive in cells

**DOI:** 10.1101/2023.02.24.529920

**Authors:** Eric A. Vitriol, Melissa A. Quintanilla, Joseph J. Tidei, Lee D. Troughton, Abigail Cody, Bruno A. Cisterna, Makenzie L. Jane, Patrick W. Oakes, Jordan R. Beach

## Abstract

Directed transport of cellular components is often dependent on the processive movements of cytoskeletal motors. Myosin 2 motors predominantly engage actin filaments of opposing orientation to drive contractile events, and are therefore not traditionally viewed as processive. However, recent *in vitro* experiments with purified non-muscle myosin 2 (NM2) demonstrated myosin 2 filaments could move processively. Here, we establish processivity as a cellular property of NM2. Processive runs in central nervous system-derived CAD cells are most apparent as processive movements on bundled actin in protrusions that terminate at the leading edge. We find that processive velocities *in vivo* are consistent with *in vitro* measurements. NM2 makes these processive runs in its filamentous form against lamellipodia retrograde flow, though anterograde movement can still occur in the absence of actin dynamics. Comparing the processivity of NM2 isoforms, we find that NM2A moves slightly faster than NM2B. Finally, we demonstrate that this is not a cell-specific property, as we observe processive-like movements of NM2 in the lamella and subnuclear stress fibers of fibroblasts. Collectively, these observations further broaden NM2 functionality and the biological processes in which the already ubiquitous motor can contribute.

## Introduction

The ability for actin- and microtubule-based molecular motors to processively “walk” along tracks, typically carrying cargo attached to their tails, is critical throughout physiology (1–5). A simplified motor mechanochemical cycle includes ATP hydrolysis, track binding, *P*_*i*_ release, a power stroke, track unbinding, and nucleotide exchange (6). Processive motors are often dimers whose ability to walk along a cytoskeletal filament is dependant on the coordinated mechanochemical cycle and track binding of the motor pair to prevent the motor from diffusing away (7, 8). The fraction of each mechanochemical cycle a motor spends in its track-bound state, also known as its duty ratio, is thus critical for processivity. Most processive motors have a high duty ratio, spending the majority of their mechanochemical cycle in the track-bound state (9), though increasing the number of motors in an ensemble can enhance the synergistic processivity of low duty ratio motors (10).

Myosin 2s are low duty ratio motors (11, 12) that are the dominant contractile motor proteins in all cells (13). The myosin 2 superfamily includes striated myosin 2 (cardiac and skeletal), smooth muscle myosin 2, and non-muscle myosin 2 (NM2). Mammalian cells express up to three isoforms of non-muscle myosin 2 (NM2A, NM2B, and NM2C) which are derived from three distinct genes (*MYH9, MYH10*, and *MYH14*, respectively) (14–16). All myosin 2s form hexameric “monomers” that consist of two myosin heavy chains (MHC), two essential light chains (ELC), and two regulatory light chains (RLC) (17). This holoenzyme is referred to as a monomer because they dynamically polymerize, or assemble, into bipolar filaments with motor domains at opposing ends. Myosin 2 filaments contain between *∼*15-300 monomers depending on the isoform (18–20). While NM2 monomers are generally considered non-processive, NM2 filaments that contain many monomers could remain track-bound for extended periods because of the high probability that at least one of its motor heads is in contact with the actin filament at any given time. Indeed, NM2A and NM2B filament processivity has been observed *in vitro* (21), though this required the presence of crowding agents for NM2A. No cellular observations or characterization for NM2 processivity, to our knowledge, has been reported.

The lack of apparent NM2 processivity in cells could be attributed to the architecture of most actin networks. If the bipolar filament is presented actin filaments of opposing orientation, then both sets of motors can engage the respective actin, hydrolyze ATP, and produce force. This is the basic mechanism of contractile force generation that drives muscle contraction (22, 23), cell division (24), and many more myosin 2-dependent processes. However, if a bipolar filament is presented with a single actin filament, or a parallel actin bundle, then one set of motors can engage the actin while the opposing motors on the opposite end of the filament are less likely to engage. In this case, the engaging motors can dominate and processive movements are possible. Muscle myosin 2s, especially striated, are unlikely to encounter parallel actin bundles, as the sarcomere is vitally assembled with individual actin “thin filaments” in opposing orientation to drive contraction (25). However, because NM2s are expressed in such a diverse array of cell types, they are more likely to encounter parallel actin bundles (26) that could support processive movements. Here, we provide the first cellular observations and characterization of myosin 2 processivity. Using multiple tagging approaches, we demonstrate processive filamentous NM2 that is independent of actin dynamics but is dependent on actin architecture. We also observe subtle kinetic differences between NM2 isoforms consistent with previous *in vitro* experiments. Finally, we show that processive-like movements occur in multiple cell types, demonstrating that this is not a cell-specific NM2 feature.

## Results

### Tagging RLC reveals NM2 Processive-like Anterograde Movements in CAD cells

Cath.a-differentiated (CAD) cells are a central nervous system derived cell line (27) that form broad actin-based lamellar protrusions, reminiscent of the leading edge of a neuronal growth cone. These lamellae contain both Arp-dependent actin mesh networks and Mena/VASP-dependent linear parallel actin bundles (28). To observe all NM2 localization and dynamics in these protrusions, we expressed RLC with a C-terminal 3x-iRFP670 tag (RLC-iRFP; Fig. 1A). We observed filaments throughout the cell body, along transverse arcs, and nascent filament clusters in the protrusion (Fig. 1A). As expected, many of these NM2 clusters moved retrograde towards the cell center, coupled with actin retrograde flow (Fig. 1B - 1E, blue indicators; Supplemental Movie S1). However, unexpectedly, we also observed significant NM2 anterograde movements toward the leading edge (Fig. 1B - 1E, orange indicators). These anterograde-moving puncta mostly originated from dense regions in the posterior protrusion, though they also occasionally partitioned off of existing NM2 filaments (Fig. 1C, green arrows; Supplemental Movie S2). Anterograde puncta often reached the leading edge as in Fig. 1C. Using automated tracking analysis, we measured the kinetics of these retrograde and anterograde movements, finding retrograde velocities of *∼*35 nm/s in magnitude (Fig.1E, blue) and anterograde velocities of *∼*60 nm/s (Fig. 1E, orange). These values are consistent with actin retrograde movements in these cells (28, 29), and on a similar scale to *in vitro* processive measurements for NM2A and NM2B (*∼*120nm/sec and *∼*40 nm/sec, respectively)(21), especially when accounting for retrograde flow.

**Fig. 1.**
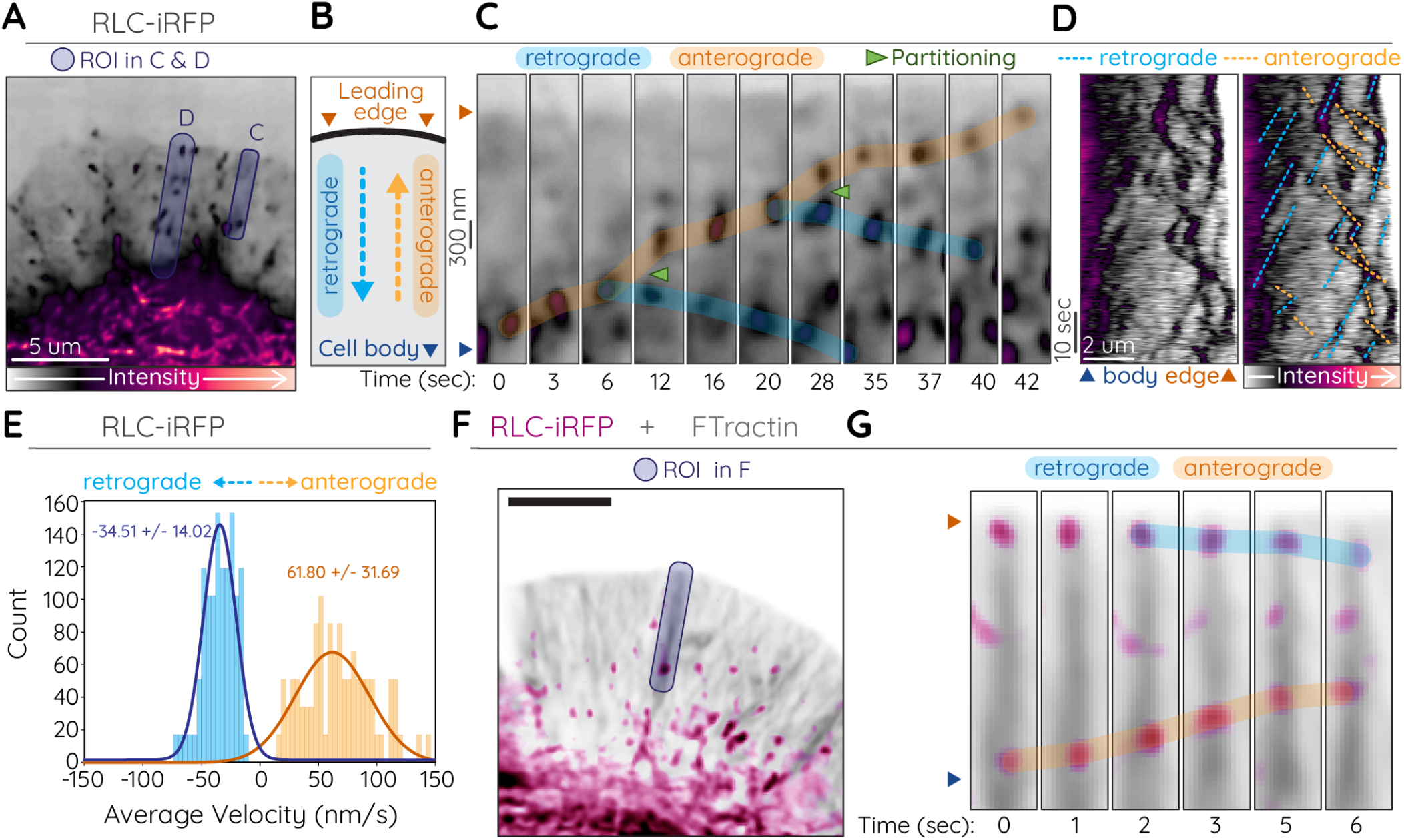
Processive-like anterograde movements by RLC. (A-E) CAD cell expressing RLC-iRFP. Purple translucent regions in (A) indicate ROI for (C) and (D). B) Cartoon depicting location of cell body and leading edge and direction of retrograde (dotted blue arrow) and anterograde (dotted orange arrow) movements. C) Time-lapse of anterograde (orange highlight) and retrograde (blue highlight) motion of RLC puncta. Partitioning filaments indicated by green arrows. D) Duplicate kymographs with (right) or without (left) retrograde and anterograde movements indicated. E) Histograms of velocity measurements from automated tracking of RLC-iRFP retrograde and anterograde movements in protrusions (see Table 1). F and G) CAD cell expressing RLC-iRFP (magenta) and FTractin-mScarlet (grey). Blue translucent ROI in (F) indicates region used for time-lapse in (G). G) Selected time points demonstrating retrograde (blue highlight) and anterograde (orange highlight) motion along an actin bundle.

To determine the underlying actin architecture on which these anterograde movements were occurring, we co-expressed RLC-iRFP with the filamentous actin reporter FTractin tagged with mNeon-Green (FTractin-Neon; Fig. 1F-G, Supplemental Movies S3-4). We readily observed anterograde RLC movements (magenta) on thick actin bundles (grey) that terminate at the leading edge. This architecture is consistent with parallel actin bundles assembled by the Mena/VASP family (28).

### Tagging MHC reveals NM2 Processive-like Anterograde Movements in CAD cells

While these observations are consistent with processive NM2, RLCs are promiscuous and can bind other members of the myosin superfamily (30–33). We sought, therefore, to confirm that these anterograde movements were indeed filamentous NM2 movements by expressing NM2A with an N-terminal EGFP (EGFP-NM2A). Similar to the RLC, we observed filamentous structures in the cell body with nascent filaments appearing in the protrusion (Fig. 2A). We again observed both retrograde and anterograde movements (Fig. 2B-C; Supplemental Movie S5), and observed filaments partitioning while moving anterograde (Fig. 2B, green arrows), before eventually reaching the leading edge or stalling and moving retrograde.

**Fig. 2.**
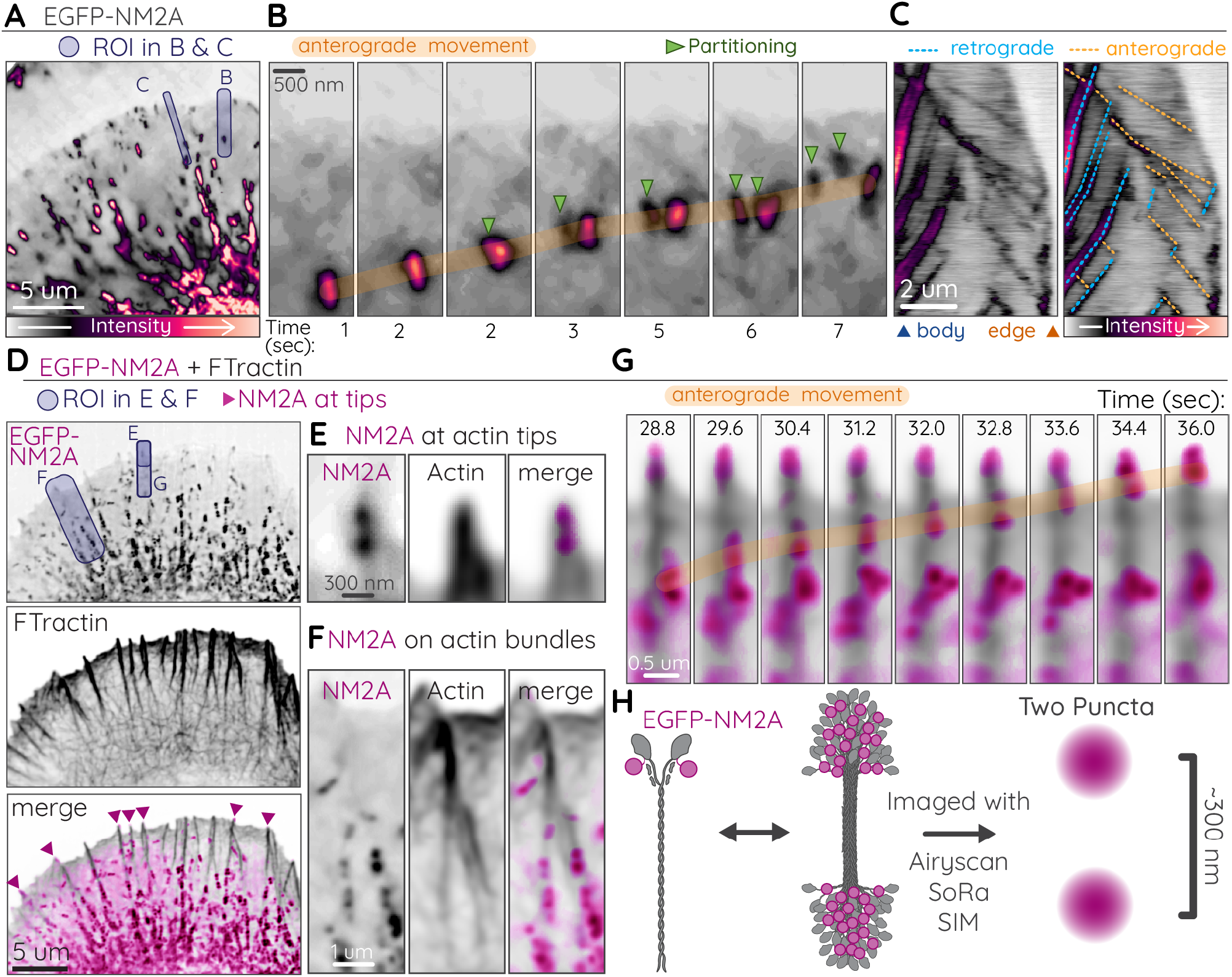
Processive-like Anterograde Movements by Filamentous NM2A. A-C) A representative CAD cell expressing EGFP-NM2A. Purple shaded boxes indicate ROIs for (B) and (C). B) Time-lapse of anterograde moving NM2 filament (orange highlight) that partitions twice (green arrows). C) Kymograph of EGFP-NM2A (left) with retrograde (blue dotted lines) and anterograde (orange dotted lines) movements indicated (right). Blue arrow and orange arrow below left kymograph indicate cell body and leading edge, respectively. D) CAD cell expressing EGFP-NM2A (magenta) and FTractin-mScarlet (grey). Purple shaded boxes indicate ROIs for (E), (F) and (G). E) NM2A filament clearly localizing at a protrusive actin tip. F) NM2A filaments accumulating along actin bundles. G) Time-lapse anterograde movements of NM2A (orange highlight). H) Cartoon of N-terminal EGFP tag on NM2A monomer, filament, and filament imaged with high resolution microscopy.

Previous EM studies have observed activated monomeric NM2 in cells (34), and NM2 is known to associate with various vesicular organelles that undergo directed transport (35–38). To confirm that our anterograde movements were filamentous NM2 structures moving on actin, we imaged both EGFP-NM2A and FTractin-mScarlet in CAD protrusions (Fig. 2D-F; Supplemental Movie S6). N-terminal tagging of NM2A coupled with high-resolution imaging reveals two puncta *∼*300 nm apart, indicative of bipolar filamentous NM2 (Fig 2H)(39, 40). We readily observed these bipolar filaments at the tips of short actin protrusions (Fig. 2E), on actin bundles (Fig. 2F), and moving anterograde along actin bundles (Fig. 2G). Collectively these different tagging approaches of the NM2 holoenzyme demonstrate the anterograde movements of filamentous NM2 along actin bundles near the leading edge.

### NM2 moves anterograde independent of actin dynamics

We next sought to isolate movement of the NM2 filaments from the underlying actin dynamics. We used a small-molecule inhibitor cocktail of jasplakinolide and latrunculin (JL) to stall actin dynamics, similar to previous work (41), and monitored NM2 by expressing NM2A with a C-terminal HaloTag (NM2A-Halo). This places all fluorophores into a single punctum at the middle of the bipolar filament (Fig. 3A), in contrast to N-terminal tags which creates two puncta at opposing ends of the bipolar filament (Fig. 2H). This C-terminal HaloTag provides higher signal to noise to closely monitor filament dynamics via kymographs or automated tracking. Prior to JL treatment, we observed both retrograde and anterograde movements of NM2A-Halo (Fig. 3B-C,F; Supplemental Movie S7). After JL treatment, NM2A-Halo retrograde flow was largely abrogated, as evidenced by the vertical lines in the kymograph and reduced movements detected in automated tracking (Fig. 3D-F). This is consistent with NM2 retrograde movements being dependent on actin dynamics. After JL treatment, however, we still observed robust anterograde movements (Fig. 3D-F; Supplemental Movie S7). In addition, inhibition of retrograde flow increased the leading edge localization of NM2 (Fig. 3D,G). Collectively, these data illustrate that the anterograde movements we observe are processive NM2 filaments and that NM2 can move processively in the absence of actin dynamics.

**Fig. 3.**
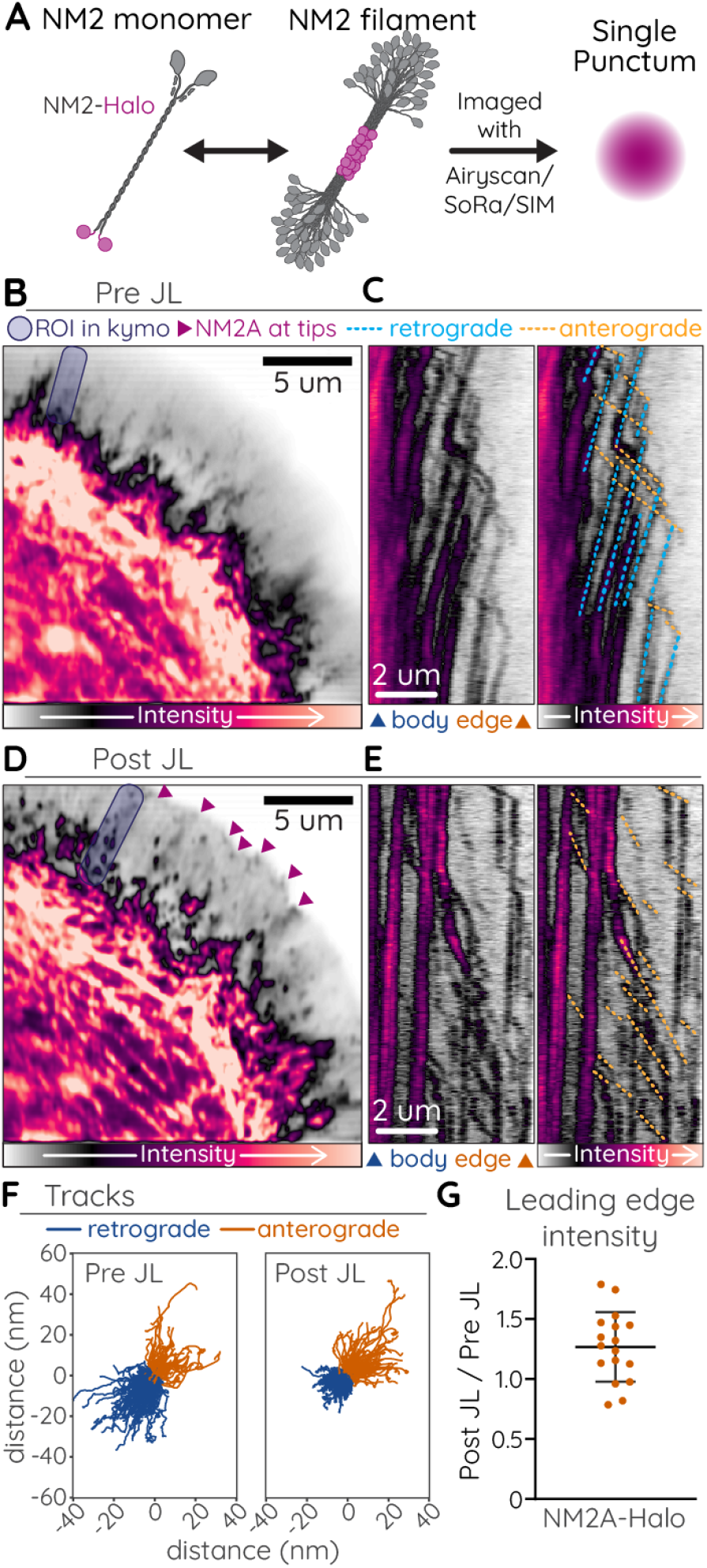
NM2 anterograde movements independent of actin dynamics. A) Cartoon of C-terminal HaloTag on an NM2 monomer, filament, and imaged with high resolution microscopy. B and D) CAD cell expressing NM2A-Halo imaged before (B) or after treatment with jasplakinolide/latrunculin (JL) cocktail. Purple shaded boxes indicate ROIs for (C) and (E). Magenta arrows indicate NM2 at apparent tips of leading edge actin bundles. C and E) Kymographs shown with (right) and with (left) retrograde (blue dotted lines) and anterograde (orange dotted lines) movements indicated. F) Rose plot of individual retrograde (blue) and anterograde (orange) tracks detected in a single cell before and after JL treatment. G) Ratio of NM2A-Halo intensity within 1 um of the leading edge 45 seconds post-JL treatment relative to last time point before treatment.

### Processive NM2 is dependent on actin architecture

Previous studies in CAD cells demonstrated that knocking out the monomeric actin binding protein profilin 1 (PFN1 KO) disrupts lamellipodia and pre-filopodial actin bundling, while over-expression of profilin enhances actin bundles at the expense of Arp2/3-dendritic networks (28). We therefore used different levels of profilin 1 expression to determine if altering the lamellipodia actin network affected NM2 anterograde movements. When NM2A-Halo was expressed in PFN1 KO CAD cells, it appeared largely diffuse and non-filamentous, with little apparent retrograde flow and few discernible processive anterograde movements (Fig. 4A-B; Supplemental Movie S8). This lack of NM2 architecture and apparent decreased assembly is not surprising, considering the extent to which PFN1-KO disrupts actin structures in these cells (28). However, it does support a model that appropriate actin architecture in the form of linear arrays is required for processive NM2 anterograde runs. Indeed, anterograde moving NM2A-Halo labeled filaments were readily observed in membrane protrusions in CAD cells overexpressing PFN1 (Fig. 4E; Supplemental Movie S8). Though the number of initiated anterograde events and run velocity were no different than controls (GFP; Fig. 4G-4I), we did observe a significantly higher number of anterograde moving NM2A puncta that reached the leading edge (Fig. 4J). We also observed persistent localization of these puncta at the tips of short actin spikes, likely pre-filopodial bundles (Fig. 4C; magenta arrows). The PFN1 overexpression data suggest that dendritic actin networks inhibit the ability of NM2A to move processively along pre-filopodia actin bundles. This model could explain why NM2A anterograde movements have not been reported in lamellipodia that are more dependant on Arp2/3 networks, like those found in migratory fibroblasts.

**Fig. 4.**
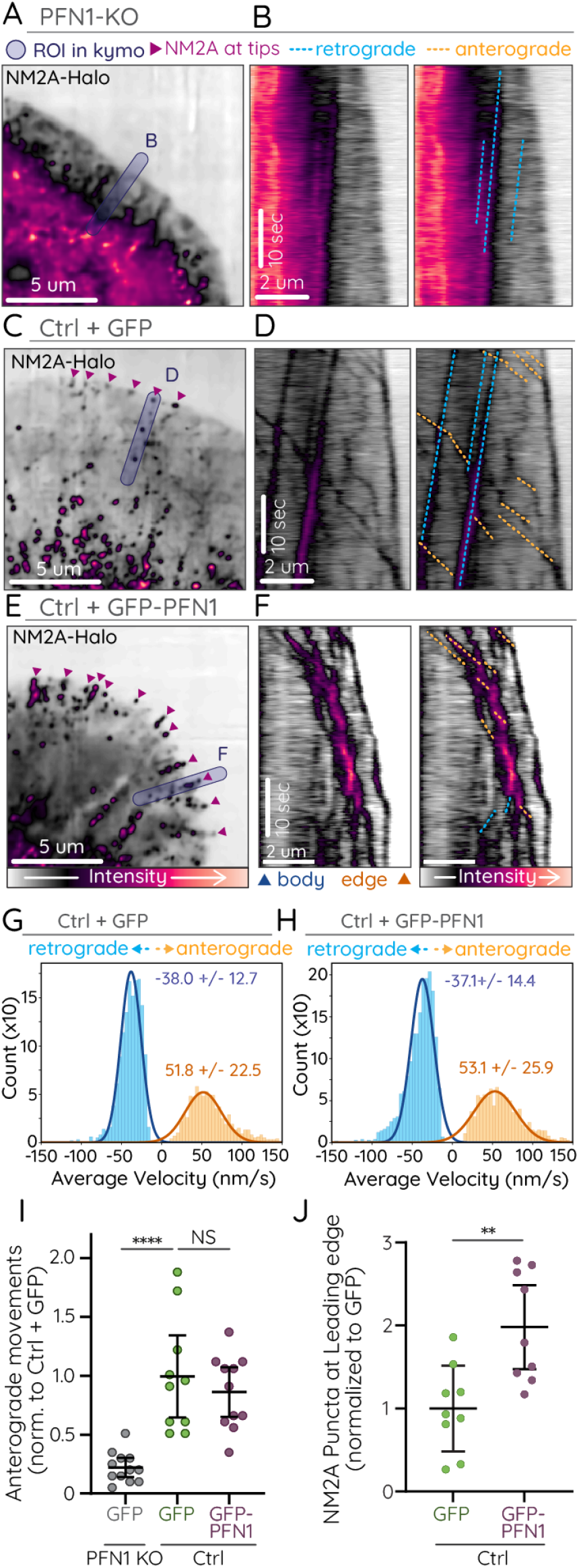
Processive NM2 is dependent on lamellipodia architecture. NM2A-Halo was imaged in PFN1-KO CAD cells, CAD cells expressing GFP, or CAD cells overexpressing GFP-PFN1. A-F) Representative images (A, C, E) and duplicate kymographs (B, D, F) with (right) and without (left) annotation of anterograde (orange dotted lines) and retrograde (blue dotted lines) NM2A movements in PFN1 KO (A,B), ctrl + GFP (C,D) and ctrl+ GFP-PFN1 (E,F) cells. G,H) Histograms of velocity measurements from automated tracking of NM2A-Halo puncta in control and PFN1 overexpressing cells (see Table 1). I) Plot of the number of NM2A-Halo anterograde movements in the lamellipodia protrusions of PFN1 KO, control, and PFN1 overexpressing cells. Only anterograde events that initiated within 8 µm of the leading edge were counted. Each dot indicates one cell. Error bars depict mean +/- SD. J) Number of NM2A-Halo puncta within 1 µm of the leading edge of PFN1 KO, control, and PFN1 overexpressing cells. Each dot indicates one cell. Error bars depict mean +/- SD.

### Characterization of NM2 Isoform Processivity

To better understand processivity of NM2 isoforms, we evaluated the endogenous RNA, protein, and localization in CAD cells. Previous RNAseq data (28) demonstrated that MHC 2A (*Myh9*) and MHC 2B (*Myh10*) were transcribed at moderate levels, with *∼*2:1 ratio of MHC 2A:MHC 2B (mean = 2.20 +/- 0.28), while MHC 2C (*Myh14*) was barely detected (Fig. 5A). Western blotting of CAD whole-cell lysates demonstrated similar expression levels of MHC 2A and MHC 2B (Fig. 5B). Correlative blots with EGFP intermediates revealed MHC 2A dominant over MHC 2B expression, again with an *∼*2:1 ratio of MHC 2A:MHC 2B (mean = 1.95 +/- 0.66). Immunostaining of CAD cells also revealed robust NM2A signal with clear puncta extending into the protrusion, while NM2B signal was less continuous throughout the cell, with less signal above background in the protrusion (Fig. 5C). This is consistent with previous localization in polarized cells, where NM2A extends more peripheral and NM2B remains more centripetal and rearward (42).

**Fig. 5.**
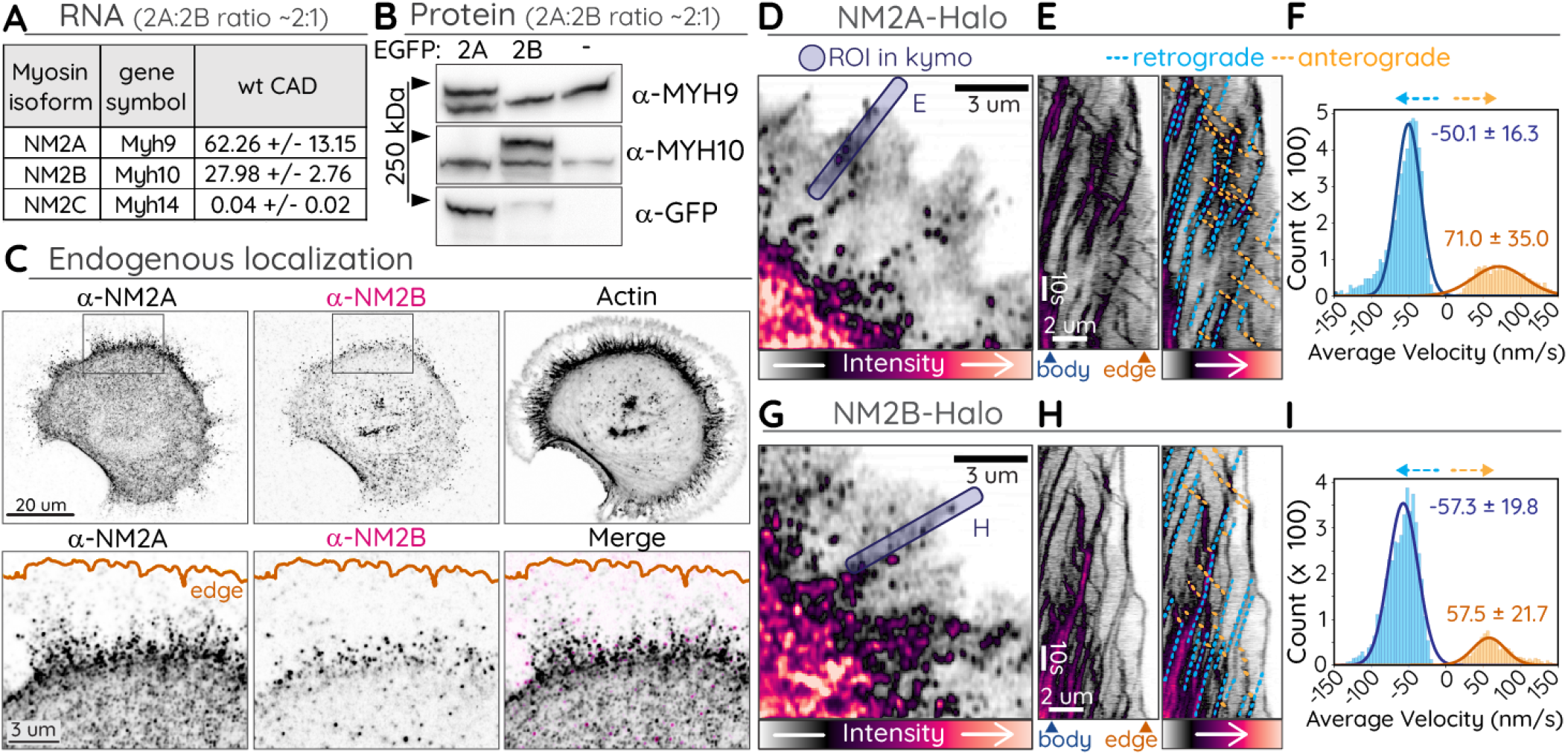
Differential expression, localization and velocity of NM2 isoforms. A) Relative RNA expression of NM2A MHC genes. B) Whole cell lysates of untransfected CAD cells or cells over-expressing EGFP-NM2A or -NM2B were subject to western blotting with the indicated antibody. The NM2A:NM2B ratio was calculated using anti-GFP blot as intermediate (see Methods). C) CAD cells fixed and immunostained with indicated NM2 isoform-specific antibody and phalloidin (F-actin). Box in top row indicates ROI for insets in bottom row, which includes merge (rightmost panel). D-I) CAD cells expressing NM2A-Halo (D-F) or NM2B-Halo (G-I). Blue boxes indicate ROI for kymographs in and (H). E and H) Kymographs shown with (right) and without (left) retrograde (blue dotted lines) and anterograde (orange dotted lines) movements indicated. F and I) Histograms of velocity measurements from automated tracking of NM2A-Halo and NM2B-Halo retrograde and anterograde movements in protrusions (see Table 1).

In addition to localization differences, NM2 isoforms were found to possess kinetic differences in processivity *in vitro* (21). To determine if we could detect any differences in processivity kinetics in cells, we expressed NM2A-Halo or NM2B with a C-terminal HaloTag (NM2B-Halo). When NM2A-Halo was expressed in CAD cells, we again observed robust processivity in the protrusions, with many runs reaching the leading edge (Fig. 5D-E; Supplemental Movie S9). When NM2B-Halo was expressed in CAD cells, we observed processive runs but with clear differences relative to NM2A. NM2B runs again initiated from the dense cell body or from partitioning events off of retrograde moving NM2 clusters. However, the runs often stalled in the mid-protrusion region before reaching the leading edge (Fig. 5G-H). Average velocity measurements revealed a mild but significant difference between the isoforms, with NM2A running over *∼*70 nm/sec and NM2B running under *∼*60 nm/sec (Fig. 5F and 5I; Table S1).

### Processive NM2 in Fibroblasts

While the actin architecture and morphology of CAD cells provide opportunity for readily observable processive NM2 movements, we hypothesized that similar movements could and should still be occurring in diverse cell types, if perhaps less frequently. To explore this possibility, we imaged primary mouse embryonic fibroblast (MEF) cells with NM2A endogenously labeled with EGFP (EGFP-NM2A). When sampled with high spatial and temporal frequency, we observed processive NM2 movements in both subnuclear stress fibers (Fig. 6A-B; Supplemental Movie S10) and ventral stress fibers in the lamellar protrusions (Fig. 6C-D; Supplemental Movie S11).

**Fig. 6.**
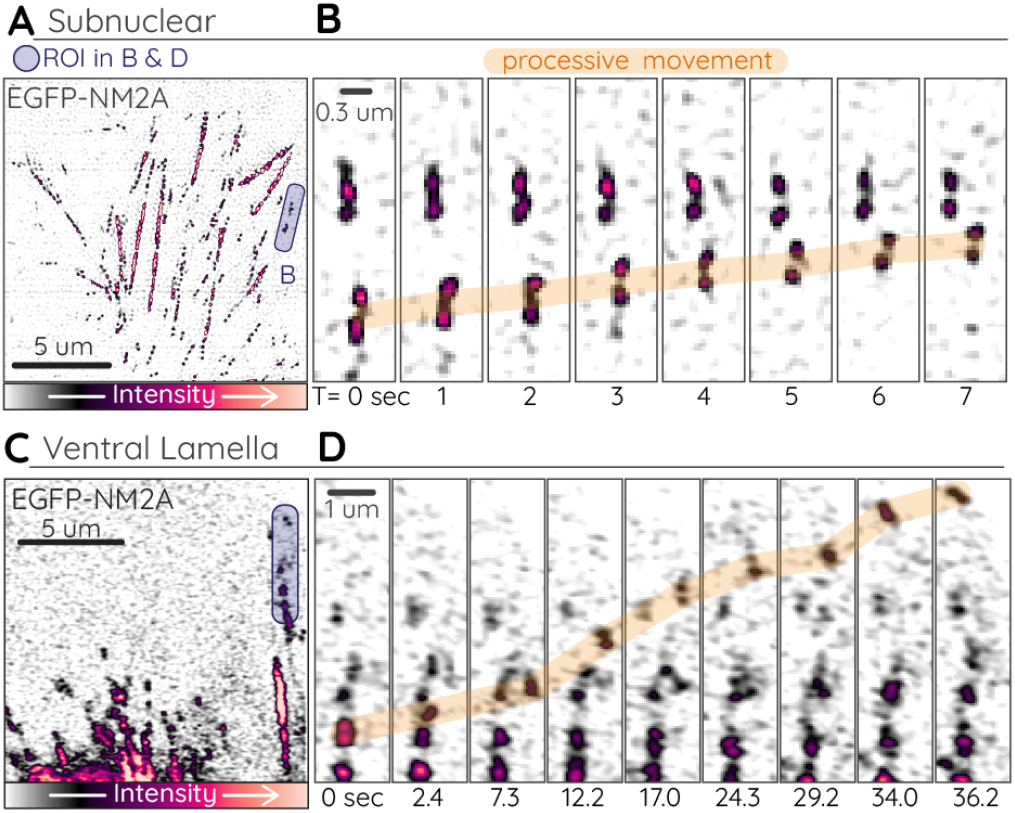
Processive EGFP-NM2A in fibroblasts. A-D) Primary EGFP-NM2A MEF cells imaged with total internal reflection fluorescence structured-illumination microscopy (subnuclear stress fibers; A and B) or Airyscan (ventral stress fibers; C and D). Purple boxes in (A) and (C) indicate ROI for time-lapses in (B) and (D), where processive movements are indicated with orange highlight.

## Discussion

We conclude that the anterograde movements we observe for NM2 are indeed processive for the following reasons: first, they travel against the direction of actin retrograde flow; second, the velocity of movement is consistent with *in vitro* measurements of NM2 processivity (21); third, they often travel along actin bundles that terminate at the leading edge, the exact actin architecture where NM2 processivity would theoretically thrive; and finally, stalling actin dynamics did not inhibit movement. These results, in combination with the previous *in vitro* work by Sellers and colleagues, establishes processivity as a property of NM2 filaments. Our collective observations provoke a range of questions, from the biophysical nature to the biological relevance of processive filaments.

Previous studies have used a simple formula to estimate the likelihood of synergistic processivity for an ensemble of motors:

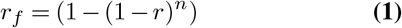

where *r*_*f*_ is the duty ratio of a filamentous ensemble, *r* is the duty ratio of an individual motor, and *n* is the number of motors in a filamentous ensemble (21, 43). As *r*_*f*_ increases, the likelihood of processivity increases. Melli et al. calculated NM2A could be processive with a duty ratio of 0.05 and *∼*50 motors per side of the bipolar filament (*∼*25 “monomers” per side with two motors per monomer) (21). Considering NM2A filaments mature with *∼*30 monomers per filament (*∼*30 motors per side) (18), only two mature filaments would be required to create a processive ensemble. Previous EM studies (44, 45) demonstrate the existence of filament stacks in the lamella, with multiple filaments in register. If many lamellar filamentous structures are indeed small sub-resolution stacks with at least two filaments, this would provide sufficient motors to maintain binding to the actin track and enable NM2A to move processively towards the leading edge.

A similar theoretical approach was taken by Nagy et al. who calculated that NM2B would be processive with a duty ratio of 0.22 and *∼*10-12 motors per side of the bipolar filament (43), although previous contrasting work suggested NM2B monomer processivity (46). This theoretical 10-12 NM2B motors is easily attained in a single filament. However, the increased duty ratio of NM2B relative to NM2A should lead to more motors engaged with filamentous actin, especially if there are multiple mature filaments in an ensemble in a dense actin network. We hypothesize that this increased actin binding explains our observation that NM2B filaments often struggle to reach the leading edge, as they are more easily entangled in the lamellar actin network. In addition, the moderate decrease in NM2B velocity relative to NM2A could be explained by this increased NM2B duty ratio and decreased ATPase kinetics (47), adding drag to the ensemble with decreased motor cycling.

In reality, our kinetic measurements are complicated by a number of factors. The cellular actin architecture is more complex than the *in vitro* assays. Cellular actin is decorated with an array of actin binding proteins, unlike the pure actin filaments *in vitro*. Any of these actin binding proteins could potentially enhance or inhibit processive NM2. Cellular NM2 filaments are in dynamic equilibrium with monomeric NM2, and could be bound to numerous binding partners (see below). Perhaps most importantly, NM2 isoforms have the ability to co-assemble into mixed filaments (34, 48). The cell types analyzed in this study are skewed towards NM2A expression over NM2B. Our immunostaining of endogenous isoforms, consistent with previous observations (42), demonstrates spatial sorting of NM2 filaments, with NM2B restricted to central regions in the cell body and NM2A extending more peripherally. Therefore, our kinetic measurements of peripheral EGFP-NM2A processivity are likely to be measuring filaments largely composed of NM2A, while EGFP-NM2B processive filaments in more centripetal regions are likely to be mixed NM2A/NM2B filaments. When NM2A and NM2B were mixed 2:1 *in vitro* and allowed to co-polymerize, the velocity was indistinguishable from NM2B alone (21), suggesting that relatively small numbers of NM2B motors can significantly alter filamentous biophysical and kinetic properties. Therefore, we hypothesize that the EGFP-NM2B kinetics we measure for mixed NM2A/NM2B filaments are revealing the underlying properties of NM2B processivity. More sophisticated experimental approaches are required to further dissect these concepts. A curious question is “why has NM2 processivity not been previously reported?” Certainly, there has been an abundance of live-cell imaging of NM2. We suspect a confluence of factors can explain this discrepancy. Both subcellular conditions and experimental approach must be optimal. First, there must be a relatively high ratio between parallel actin bundles and other actin structures (antiparallel bundles, mesh networks, etc.). This excludes many stress fibers, transverse arcs, contractile rings, cortices, lamella and lamellipodia. Second, actin retrograde flow must be significantly slower than the velocity of NM2 processive movements in order for the NM2 processive movements to be observable. This is especially true for NM2B, which has slower velocities *in vitro* and in cells. Considering retrograde flow in some immune cell lamellipodia can be *∼*100 nm/sec (49), many anterograde NM2 movements would be difficult to observe. Third, NM2 density must be sufficiently low to observe filament dynamics. Many regions of non-muscle cells contain dense actomyosin networks, and discerning the movements of individual filaments or small filamentous ensembles is beyond the resolution of most light microscopy. Considering these caveats, we suggest that protrusions with highly bundled parallel actin, as can be found in neuronal growth cones, filopodia, stereocilia, invadopodia, etc., are the most likely subcellular environments where NM2 processivity might be apparent. Consistently, previous immmunostaining of neuronal growth cones demonstrated ample NM2 signal in peripheral structures and filopodia (50, 51). Finally, concerning experimental approach, temporal sampling frequency needs to be sufficiently high (*∼*1-2 Hz) that processive movements can be revealed. Studies that observe NM2 dynamics in slower cellular processes (e.g. migration, cytokinesis, epithelial dynamics, etc.) by sampling with longer intervals are likely to miss any processive movements.

The most outstanding question remains as to the biological function of processive NM2. One obvious answer is that as a processive motor NM2 could act as a cargo transporter. Numerous NM2 binding partners have been identified (52–55), and NM2 has been localized to various vesicle and organelle populations (exocytic vesicles, lytic granules, lipid droplets, Golgi vesicles, etc.) (35–38). It is possible that these or other cargos are being transported to the distal regions of these parallel actin structures. However, as other specialized processive myosins (e.g. myosins V, X and XV (56, 57)) exist with this capability, it is unclear why the cell would rely on NM2 for this function. One also wonders if processive NM2 in these structures would compete with or complement the function of these other myosins. Another potential model for the functional relevance of NM2 processivity is that it serves to localize NM2 itself into peripheral structures with parallel actin bundles. This model, however, seems inefficient compared to the numerous phosphorylation assembly mechanisms that are used throughout cell biology (52, 58). In addition, signaling pathways exist that specifically inhibit NM2 assembly at the leading edge (59). It would be interesting to test if these pathways are turned off in CAD cells or during PFN1 over-expression. Manipulation of NM2 to dissect these models is likely to prove challenging. Studies that broadly inhibit NM2 function throughout the cell via knockdown/KO or small-molecule inhibitors (e.g. blebbistatin) would be difficult to interpret, as they would disrupt the entirety of the actomyosin network. Photomanipulation approaches that can target NM2 specifically in protrusions (e.g. iLID-based inhibitor (60)) might provide the spatio-temporal precision required. Finally, correlative imaging with NM2 and potential cargos, could also prove insightful but requires identifying potential cargos to make it practical.

In conclusion, the identification of NM2 processivity forces us to broaden the potential functional roles for NM2 in cell physiology. While its dominant function is to drive contractility, its ability to function as a monomer, to drive actin expansion, to cross-link actin, and to move processively should not be overlooked.

## Materials and Methods

### Cell culture and transfection

Cath.-a-differentiated (CAD) cells were purchased from Sigma-Aldrich and cultured in DMEM/F12 medium (Gibco) supplemented with 8% fetal calf serum, 1% L-Glutamine, and 1% penicillin-streptomycin. Two to four hours prior to imaging, CAD cells were plated on coverslips coated overnight at 4 °C with 10 µg/mL laminin (Sigma-Aldrich). DMEM/F12 medium without phenol red (Gibco) supplemented with 15mM HEPES was used for live-cell imaging. Cell lines were also routinely tested for mycoplasma using the Universal Detection Kit (ATCC). PFN1 KO cells were generated with CRISPR/Cas9 as previously described (28). CAD cells were transfected with plasmid DNA via electroporation as previously described (28) or with LipoD293 (SignaGen, “Hard-To-Transfect Mammalian Cell” protocol). Cells expressing HaloTag constructs were incubated overnight with 10-100 nM Janelia Fluor 646 HaloTag ligand (Promega) (61). EGFP-NM2A knock-in MEFs were generated from mice (62), and isolated and cultured as previously described (40).

### Plasmids

NM2A-Halo (pHalo-N1-NM2A) was generated by swapping HaloTag for mApple in pmApple-N1-NM2A, which was previously described (48). NM2B-Halo (pHalo-N1-NM2B) previously described (40). EGFP-NM2A was a gift from Dr. Thomas Egelhoff and is available at Addgene (https://www.addgene.org/11347/). RLC-iRFP was generated by removing FTractin from pLV-3x-iRFP670-FTractin (gift from Dr. John A. Hammer, NHLBI/NIH, Bethesda, MD) and introducing a gBlock (IDT) containing RLC coding sequence using Gibson cloning. FTractin-EGFP was a gift from M. Schell (Uniformed Services University, Bethesda, MD) (63). FTractin-mApple was previously described (48).

### Antibodies

NMIIA (a.a.1936-1950, ECM Biosciences) Rabbit polyclonal antibody used at 1:2000

NMIIB (clone A-3, SC-3769-42, Santa Cruz Biotechnology) Mouse monoclonal antibody used at 1:2000

GFP (clone B-2, SC-9996, Santa Cruz Biotechnology) Mouse monoclonal antibody used at 1:2000

### Imaging

NM2 isoform analyses (immunostaining and NM2A-Halo and NM2B-Halo imaging), JL experiments, and ventral lamellar EGFP-NM2A primary MEF imaging was done on a Zeiss Airyscan 880 Microscope using a Plan-Apochromat 63x/1.4 Oil DIC M27 objective and 633 nm laser. Raw data was processed using Zen software with automated processing strength. Subnuclear stress fiber imaging in EGFP-NM2A primary MEF was done using total internal reflection fluorescence structured-illumination microscopy (TIRF-SIM), as previously described (48). To decrease the background artefact inherent in SIM processing, TIRF-SIM data were blurred once with “Smooth” in Fiji/ImageJ (64).

Imaging of CAD cells expressing RLC-iRFP, EGFP-NM2A, and NM2A-Halo in PFN1 KO/OE experiments was performed using a Nikon CSU-W1 SoRa spinning disk confocal microscope using a 100X 1.49NA SR objective and a Hamamatsu Fusion BT Camera. All images from these data sets were acquired in SoRa mode with the 2.8X magnifier. Movies used for NM2-Halo tracking were acquired with 2×2 binning to enhance signal to noise and increase acquisition speed to 5 Hz. All data from the SoRa had background noise removed using Denoise.ai (NIS-Elements, Nikon), a trained neural network that uses deep learning to estimate and remove the noise component of an image. Images acquired using 1×1 binning were also deconvolved in NIS-Elements using the Blind Deconvolution algorithm. Before analysis, images were exported into Fiji/ImageJ and corrected for photobleaching using the histogram matching algorithm.

### Western blotting

To quantify isoform ratios for NM2A and NM2B in CAD cells, whole-cell lysates were collected from untransfected, EGFP-NM2A expressing, and EGFP-NM2B expressing cells. By separating EGFP-tagged NM2 from endogenous, we obtained a ratio between the two bands for each isoform. Using an anti-GFP blot, we obtained a ratio between EGFP-NM2A and EGFP-NM2B. The final NM2A/NM2B ratio was then obtained using these three ratios: NM2A/EGFP-NM2A : EGFP-NM2A/EGFP-NM2B : EGFP-NM2B/NM2B.

### Quantification of myosin motion

All tracking analysis was performed in python. The code and example data can be found at https://github.com/OakesLab. Briefly, images were first filtered with a 15 pixel square Laplacian of Gaussian filter with a standard deviation of 2, to emphasize the myosin filaments. Filaments were then tracked using the trackpy software package (https://github.com/soft-matter/trackpy) with the following relevant parameters: feature_size = 11 pixels, memory = 2, and separation = 3. As trackpy doesn’t use sub-pixel localization the resulting tracks were filtered using a running window average over *∼*2.5 seconds (translating to 5-8 frames depending on the imaging frequency). The resulting tracks were further filtered to only consider tracks with a path length of at least 900 nm (e.g. the length of *∼*3 myosin filaments) to ensure that we were only considering persistent motion. While tracks shorter than this certainly occurred, we restricted measurements to this range to ensure we could have confidence in the measurements of the particle velocities.

To determine the direction of the flow, we first defined the image intensity center of mass. A vector drawn from the center of the image to the center of mass determined the mean direction of retrograde flow. The direction of the myosin movement was defined as the vector drawn from the first position in the smoothed track to the last. By calculating the dot product of these two vectors we were able to define an angle, *θ*, relative to the mean direction of flow. We considered any angle *θ < π/*3 to be retrograde flow, while any angle *θ >* 2*π/*3 to be anterograde flow (essentially creating 120 degree cone in each direction). Tracks with angles between these two directions were not considered in our analysis, as they were predominantly moving parallel to the edge of the cell.

Mean values for both retrograde and anterograde flow rates were determined by fitting a gaussian to the histogram of the pooled data. The data is reported as the mean of the gaussian fit *±* the standard deviation of the gaussian curve.

### Quantification of NM2A-Halo anterograde events

The number of NM2A-Halo anterograde events in lamellipodia protrusions was quantified by drawing an 8×8 µm region of interest against the leading edge and manually counting all puncta that moved >1 µm in the anterograde direction during the duration of the movie. Three 8×8 µm regions were analyzed per movie and then averaged so that each movie recieved a single score. To quantify the density of NM2A-Halo puncta at the leading edge, all puncta within a 1 µm region of interest placed against the cell edge in lamellipodia protrusions were manually counted in frames 1, 100, and 200. The number of puncta was divided by the area that was analyzed for each region, and then all puncta density values were averaged so that each movie received a single score. All manual counting was performed by researchers who were blinded to the experimental conditions and molecules being imaged.

### Statistical analysis

Number of cells analyzed and experimental replicates were as follows:

Fig. 1E = 10 cells from 3 independent repeats.

Fig. 3G = 17 cells from 3 independent repeats.

Fig. 4G = 11 cells from 3 independent repeats.

Fig. 4H = 12 cells from 3 independent repeats.

Fig. 4I = 10+ cells for each condition from 3 independent repeats.

Fig. 4J = 9 cells for each condition from 3 independent repeats.

Fig. 5F = 21 cells from 3 independent repeats.

Fig. 5I = 12 cells from 2 independent repeats

To compare the number of anterograde NM2A movements in lamellipodia protrusions of PFN1 KO, control, and PFN1 overexpressing cells (Fig. 4I), we used one-way ANOVA followed by Tukey’s multiple comparison test. To compare the number of NM2A puncta at the leading edge of control and PFN1 overexpressing cells (Fig. 4J), we used a student’s t-test. Statistical analysis was performed using Prism (GraphPad). To compare the distributions of myosin 2A and 2B in both retrograde and anterograde flows (Fig. 5F,I), we used a Kolmogorov-Smirnov test. Briefly, the cumulative distributions functions of the distributions were calculated and compared using the stats module from scipy. p-values less than 0.05 were considered significant. The anterograde distributions were found to be significantly different.

## Data availability

The data that support the findings of this study are available upon reasonable request from the corresponding authors [Vitriol, Oakes, Beach]. All code, along with example data, used to automatically analyze myosin motion is available at https://github.com/OakesLab.

## Acknowledgements

This work is dedicated to the memory of Dr. Ken Jacobson. Research reported in this publication was supported by the Maximizing Investigators’ Research Award (MIRA) (R35) from the National Institute of General Medical Sciences (NIGMS) of the National Institutes of Health (NIH) under grant number R35GM137959 to EAV and R35GM138183 to JRB, and Graduate Research Fellowship (GRFP) from the National Science Foundation (NSF) under grant number DGE-1842190 to MAQ. PWO was not directly funded for this work but participated for fun.

## Supplementary Information

**Table 1.**
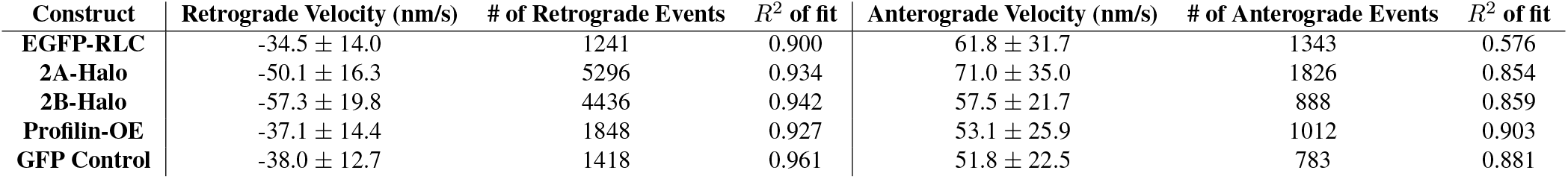

## Movie legends

**Movie S1**. CAD cell expressing RLC-iRFP from Fig. 1A. Scale bar is 5 *µ*m and time is in mm:ss. Cell was imaged at 3 Hz and playback is at 40 fps.

**Movie S2**. CAD cell expressing RLC-iRFP region of interest from Fig. 1C. Cell was imaged at 3 Hz and playback is at 40 fps.

**Movie S3**. CAD cell expressing RLC-iRFP and FTractin-Neon from Fig. 1F. Scale bar is is 5 *µ*m and time is in mm:ss. Cell was imaged at 1.5 Hz and playback is at 60 fps.

**Movie S4**. CAD cell expressing RLC-iRFP and FTractin-Neon region of interest from Fig. 1G. Cell was imaged at 1.4 Hz and playback is at 50 fps.

**Movie S5**. CAD cell expressing EGFP-NM2A from Fig. 2A. Scale bar is is 5 *µ*m and time is in mm:ss. Cell was imaged at 3 Hz and playback is at 40 fps.

**Movie S6**. CAD cell expressing EGFP-NM2A and FTractin-mApple from Fig. 2D. Scale bar is is 5 *µ*m and time is in mm:ss. Cell was imaged at 1.25 Hz and playback is at 15 fps.

**Movie S7**. CAD cell expressing NM2A-Halo treated with JL at time t=0, from Fig. 3B,D. Scale bar is is 5 *µ*m and time is in mm:ss. Cell was imaged at 2 Hz and playback is at 30 fps.

**Movie S8**. CAD cell with either PFN1-KO (left), GFP-control (middle) or GFP-PFN1 (right) expressing NM2A-Halo from Fig. 4A,C,E. Scale bar is is 5 *µ*m and time is in mm:ss. Cell was imaged at 5 Hz and playback is at 40 fps.

**Movie S9**. CAD cell expressing NM2A-Halo (top) or NM2B-Halo (bottom) from Fig. 5D,G. Scale bar is is 2 *µ*m and time is in mm:ss. Cell was imaged at 2 Hz and playback is at 40 fps.

**Movie S10**. Subnuclear stress fibers imaged in a primary EGFP-NM2A knock-in MEF from Fig. 6A. Scale bar is is 5 *µ*m and time is in mm:ss. Cell was imaged at 1 Hz and playback is at 40 fps.

**Movie S11**. Ventral lamella imaged in a primary EGFP-NM2A knock-in MEF from Fig. 6C. Scale bar is is 5 *µ*m and time is in mm:ss. Cell was imaged at 0.4 Hz and playback is at 40 fps.

